# A jump-driven self-exciting stochastic fish migration model and its fisheries applications

**DOI:** 10.1101/2023.07.13.548832

**Authors:** Hidekazu Yoshioka, Kazutoshi Yamazaki

## Abstract

We introduce a stochastic continuous-time model via a self-exciting process with jumps to describe a seasonal migration event of diadromous fish. The dynamics of the stored population at a point in a river, waiting for their upward migration, increases by the inflow from the downstream/ocean and decreases by the outflow due to their upstream migration. The inflow is assumed to occur at a constant rate until an Erlang-distributed termination time. The outflow is modeled by a self-exciting jump process to incorporate the flocking and social interactions in fish migration. Harvested cases are also studied for fisheries applications. We derive the backward Kolmogorov equations and the associated finite-difference method to compute various performance indices including the mean migration period and harvested populations. Detailed numerical and sensitivity analysis are conducted to study the spring upstream migration of the diadromous Ayu *Plecoglossus altivelis altivelis*.

## 1. Introduction

### 1.1. Problem background

Migratory fishes are major protein sources for human lives [1, 2]. They are also essential for nutrient transport and aquatic ecosystems in both local and global scales [3, 4]. However, many of them are at the risk of extinction and thus their effective conservation strategies are of great importance both ecologically and economically. To this end, accurate modeling and analysis of population dynamics of fish species are crucial as they can serve as key steps toward fish conservation strategies.

Among others, diadromous fish species are those adaptively migrating between the ocean and freshwater rivers to strategically take advantage of seasonally-varying resources. Examples include commercially important fish such as the salmonids [5], the sturgeon [6], Īnanga *Galaxias maculatus* in Australia and New Zealand [7], and the Ayu *Plecoglossus altivelis altivelis* in Japan [8]. Many diadromous fish species are endangered due to severe anthropogenetic factors, such as overfishing, release of non-native species, physical barriers like dams and weirs, and environmental pollution [9–11].

There are several complex and mutually interacting factors, that can trigger fish migration, including but are not limited to, the water flow discharge [12], water temperature [13], nutrient limitation [14], environmental heterogeneity or genetic predisposition [15], and social interactions [16]. It is infeasible to fully describe the fish migration mechanism due to a number of random biological phenomena that can affect both inter- and intra-migration events, as suggested in the reported data for diverse case studies [17–22]. A concise – but nonetheless sufficiently realistic – stochastic process model is therefore essential for the analysis of fish migration. Recent studies include those for the migration event in each season [23–25] as well as those on the population dynamics over several generations [26–28]. A tracking model of the long-time evolution of the diadromy [29] has also been analyzed.

Among the existing stochastic process models, those based on stochastic differential equations (SDEs) (e.g., [30]) provide with efficient analysis tools for fish migration in both modeling and applications. This is thanks to the close connection between SDEs and the associated partial differential equations, called the *Backward Kolmogorov equations* (BKEs), governing the probability laws modeled by the SDEs. Indeed, BKEs have been widely utilized for analyzing probabilistic behaviors of stochastic systems of interest in the science and engineering, such as disease transmission [31], predator-prey dynamics [32], chaotic flows [33], rare events in an active matter [34], and machine learning [35].

The existing literature includes the SDE-based study of the single migration event in the context of optimal stopping [36], differential games under environmental uncertainty [37], and optimal control [38, 39]. However, all these results are more analytical than quantitative. The developed models have not always been empirically tested against data, although their problems are motivated by real-world problems. Importantly, they do not account for the *self-exciting* nature of the fish migration pointed out in [16].

Existing self-exciting models are limited in the field. In [40, 41], they considered a related ecological mean-field model where the decision of each individual is aggregated into the macroscopic population dynamics. This is another form of the self-excitation through a feedback control law, but they do not consider fish migration. An SDE model accounting for the self-exciting nature, if it can be quantitatively verified against real data, can serve as an effective mathematical tool not only for modeling fish migration but also for fisheries management applications.

### 1.2. Objectives and contributions

Our objective is to develop an efficient SDE-based fish migration model in a river, along with its real-life applications. The model to be developed encompasses a wide range of migratory fish species such as the salmonids and sturgeon. In particular, we analyze the upstream migration of Ayu *P. altivelis* as a case study with real data. Our model is motivated by the approach developed by [16] that uses a self-exciting discrete-time stochastic process to model upstream migration. As in the case of [16], we study the population dynamics at a point in a river (such as a downstream of a weir), in terms of the inflow population, stored population, and outflow population (see Figure 1). However, different from [16], this paper develops an SDE-based continuous-time model using new self-exciting dynamics written in terms of a Poisson random measure in probability theory. The self-exciting nature is incorporated into the dynamics of the outflow population in a way that the intensity and size of the outflow depend on the stored population. The mortality of individuals in the population is incorporated using a discount factor, which effectively models an exponential population decay of the whole population.

**Figure 1.**
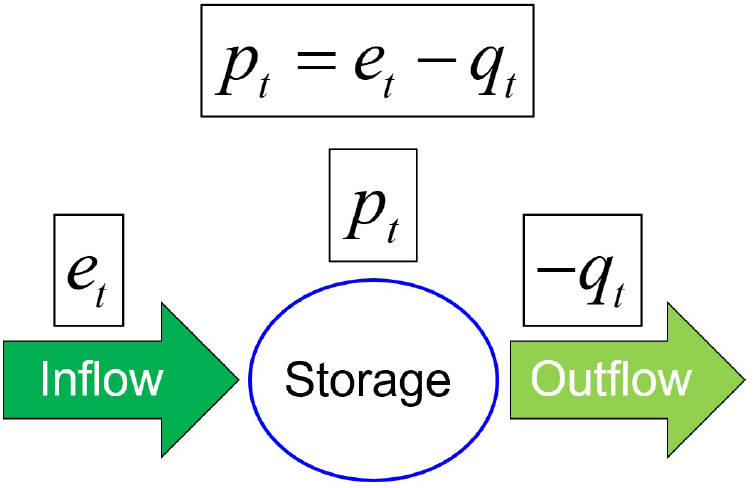
Conceptual image of the proposed stochastic process model.

The inflow is assumed to be terminated at a random time following an Erlang distribution, a generalization of the exponential distribution. We employ the Erlangization method (e.g., [42, 43]) by writing it in terms of a finite-state Markov chain, so that the proposed SDE can be formulated as a hybrid stochastic system, which is amenable to the modern stochastic calculus. A major advantage of the Erlangization method in our study is that a BKE associated with the SDE is obtained as a cascading system of partial integro-differential equations, which can be numerically discretized by using a monotone finite difference method (e.g., [44, 45]).

The BKEs along with the finite difference method are applied to the computation of the completion time of fish migration as well as the performance indices in partial harvesting problems. For the latter, we consider, what we call the *barrier-type* and *threshold-type* strategies. The barrier-type harvesting strategy can be mathematically understood as a singular control variable having a bounded variation [46, 47]. It has been applied to the long-run harvesting problem [48], while applications to the harvesting of migrating fish populations have not been addressed so far to the best of our knowledge. We also consider a more application-oriented threshold-type harvesting strategy that impulsively modulates the outflow population rather than the stored population.

We apply the proposed model to the existing record of the fish count of Ayu *P. altivelis* in a Japanese river from 2010 to 2022. We demonstrate that the computational results based on the finite difference method and a Monte-Carlo simulation agree well. Partial harvesting problems of the fish population to sustain the fish population in a river system are also investigated. We conduct detailed sensitivity analysis and study how the change of each parameter has an impact on the termination time of fish migration and harvested populations. Thus, this paper contributes to the formulation, analysis, computation, and applications of the new stochastic process model for fish migration.

The rest of this paper is organized as follows. Section 2 introduces our stochastic process model. Section 3 presents the BKEs associated with the stochastic process model, along with the finite difference scheme to discretize the BKEs. The proposed stochastic process model is applied to the real data in Section 4. The summary and future perspectives of our study are presented in Section 5.

## 2. Stochastic process model

In this section, we introduce a stochastic process model written in terms of a jump-driven SDE. We are interested in modeling the fish population dynamics at a certain location (which we call the storage) such as the downstream side of a weir in a river (see Figure 2) where the population can be measured using traps or video monitoring [12, 16, 49]. This paper studies a continuous-time model.

**Figure 2.**
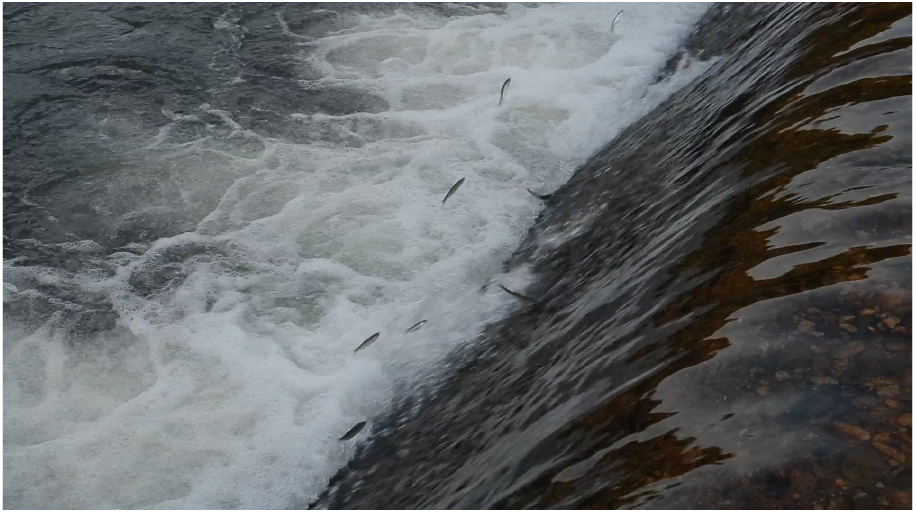
A photo of *P. altivelis* trying to jump over a weir. Location: the Hii River, Japan (supplied by the Hii River Fishery Cooperative).

On a probability space (Ω, ℱ, ℙ), we denote by *p* = (*p*_*t*_; *t* ≥ 0) the stored population at time *t*. In addition, let *e* = (*e*_*t*_; *t* ≥ 0) and *q* = (*q*_*t*_; *t* ≥ 0) be the aggregate inflow and outflow populations, respectively, until time *t*. The inflow population is for those ascending from the downstream sea or lake into the storage. The outflow population is for those migrating from the storage to the upstream side of the weir. The stored population represents those remaining in the downstream side of the weir. This formulation is qualitatively in line with [16]. For analytical tractability, we assume these processes take any non-negative values, not restricted to integers. We assume the mass conservation law that the change of the storage population is precisely determined by its inflow and outflow:

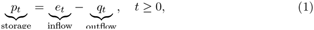

(see Remark 1 regarding mortality).

Because *e* and *q* model cumulative values, they are non-increasing processes. More details on the dynamics of the inflow and outflow populations are provided in Sections 2.1 and 2.2, respectively.

### 2.1. Inflow population

We assume a deterministic constant inflow rate *E >* 0 during a period [0, *τ*] for some random time *τ >* 0 (defined below). The inflow process *e* = (*e*_*t*_; *t* ≥ 0) starts at zero (i.e. *e*_0_ = 0) and increases at rate *E* during the inflow duration, i.e.,

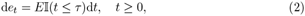

where 𝕀 (*S*) is the indicator function for a generic statement *S* such that 𝕀 (*S*) = 1 if *S* is true and 𝕀 (*S*) = 0 otherwise.

We assume that the length of the inflow period *τ* follows an Erlang distribution with shape parameter *M* and scale parameter *µ*. In other words, it is the distribution of the sum of *M* independent exponential random variables with parameter *µ*. Our choice of *τ* allows us to achieve a flexible but nonetheless analytical approach. It can be written as an absorption time of a continuous-time Markov chain *α* = (*α*_*t*_; *t* ≥ 0) on ℳ_0_:= ℳ ∪ {0} consisting of transient states ℳ:= {1, 2, …, *M*} and a single absorbing state 0 (e.g., [50, Section 3.2] and [51, Section 2]). The Markov chain *α* starts at the state *M* and, for *m* ∈ ℳ, transits from *m* to *m* − 1 at rate *µ*. More precisely,

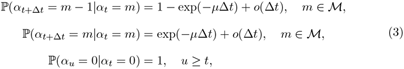

starting from the initial condition *α*_0_ = *M*. Here, *o* is the Landau symbol such that 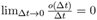. In terms of this Markov chain, we can write

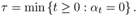

### 2.2. Outflow population

It is essential to incorporate flocking and social interactions in fish migration. Further, the existing data and survey on the outflow population size suggests that it is highly random (see e.g., [49, 52]). We thus introduce a self-exciting stochastic process to incorporate these. The aggregate outflow population process *q* is a non-decreasing pure-jump process (with right-continuous paths and left limits)

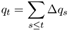

where

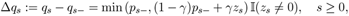

(with the understanding that *p*_0*−*_ = *q*_0*−*_ = 0) for a deterministic constant *γ* ∈ [0, 1] and the [0, ∞)-valued process *z* = (*z*_*t*_; *t* ≥ 0) described below. The jump size depends on the storage population *p*. This can be seen as a type of autoregressive (AR) process of order 1 where the parameter *γ* modulates the memory strength in the outflow population in a way that the memory persists more in the dynamics with a smaller *γ*. Capping the outflow with the current storage population *p*_*·−*_ is necessary to prevent *p* from attaining negative values (see (8)).

The process *z* determines the times and size of the jumps in *q*. It takes positive values at finitely many times on any finite interval and takes zero otherwise. At each time *t* ≥ 0, the remaining time until the next jump of *z* (i.e. forward recurrence time) is the first arrival time of a non-stationary Poisson process with rate *a* + *bp*_*·−*_ for some *a >* 0 and *b* ≥ 0. When *z*_*t*_ *>* 0, its size is an independent positive random variable with density *ν*(d·) on (0, ∞). As a result, the process *q* jumps by min (*p*_*t−*_, (1 − *γ*)*p*_*t−*_ + *γz*_*t*_). By repeating this, the aggregate outflow process *q* can be explicitly constructed.

These outflow dynamics can be modeled precisely and efficiently using the Poisson random measure in the theory of stochastic processes. We can write

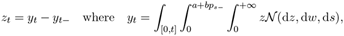

and *q*_*t*_ =∫_[0,*t*]_ d*q*_*s*_ with

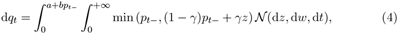

where 𝒩 is a Poisson random measure on (0, ∞)^3^ with mean *ν*(d*z*)d*w*d*t* (here d*w* and d*t* are Lebesgue measures used for jump frequency and time, respectively). For any Borel set *A* ⊂ (0, ∞)^3^, we have 𝒩 (*A*) ∼ *Poisson*(∫_*A*_ *ν*(d*z*)d*w*d*t*). In particular, for the case *b* = 0, the process *y* becomes a compound Poisson process with arrival rate *a* and jump size given by *ν*. For more details on the Poisson random measure, we refer the reader to Çinlar [53, Chapter VI].

The dynamics (4) assumes that the frequency of outflow migration depends affinely on the population size in the storage, which changes over time. The parameters *a* and *b* represent, respectively, exogenic and endogenic factors triggering the upstream migration of the stored population. The former phenomenologically corresponds to the physical disturbance such as high discharge above certain threshold values that trigger rheotaxis of the fish [12, 54]. By contrast, the latter represents the purely self-exciting effect due to flocking and social interactions of fish as discussed in [16].

### 2.3. Storage and the population dynamics

Assembling (1), (2), and (4) yields, for *t* ≥ 0,

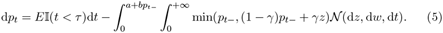

Hereafter, the stored population *p* is simply called population whenever there is no confusion. The jumps in *p* are precisely those in *q*: we define the set of outflow migration times by

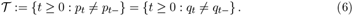

Alternatively, we can model the population dynamics in terms of a bivariate process (*p, α*) = (*p*_*t*_, *α*_*t*_; *t* ≥ 0) by keeping track of the state transition of the Markov chain describing the Erlang-distributed *τ* (defined in Section 2.1) and write, for *t* ≥ 0,

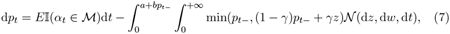

coupled with (3). It is easy to see that (*p, α*) is a time-homogeneous Markov process on ℳ_0_ × [0, ∞). The temporal evolution of the state variables (*α, p*) are determined by the above and the initial conditions (*α*_0_, *p*_0_) = (*M*, 0), which holds because at the onset no population is stored at the target site, or the downstream reach of the weir.

For each outflow migration time *t* ∈ 𝒯 (see (6)),

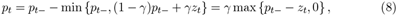

which is non-negative. This shows, together with induction arguments, that *p* is almost surely (a.s.) non-negative globally in time.

## 3. Key Performance Indices and Backward Kolmogorov equations

In this section, we introduce the key statistics and other performance indices to be analyzed in this paper and their governing BKEs. We also present a finite difference method to solve the BKEs.

### 3.1. Target statistics

#### 3.1.1. Completion time

The first target random variable we consider is the completion time of the fish migration, i.e.,

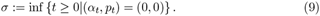

For the study of the migration of *P. altivelis*, which we will focus on later in this paper, empirical data concerning the completion time at a river in recent years is available to be used for estimating model parameter values (see Section 4). The mean completion time conditioned on (*α*_0_, *p*_0_) = (*m, x*) is denoted

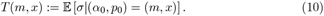

We are particularly interested in *T* (*M*, 0) because initially (at which *α*_0_ = *M*) no fish exists in the storage. We are also interested in other statistics including higher-moments, variance and standard deviation. In particular, the second-moment and the standard deviation of *σ* are denoted, respectively,

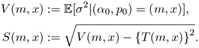

#### 3.1.2. Harvesting by a barrier-type strategy

We are also interested in fisheries application of the proposed model, in particular, harvesting. We analyze important performance indices representing harvested fish population by the following two practical strategies.

The first strategy, which we call *barrier-type harvesting strategy*, continuously replaces any excess above a prescribed level, say, *P >* 0, so that *p* always stays at or below *P*. Such control strategy is motivated by the demand for efficient measures toward the preservation/sustainability of fish population, in situations where they can become endangered due to the physical barriers such as weirs and dams that prevent them from migrating (see e.g., [44]). Harvesting the migrating fish population and distributing them to a different point in the same river system is known to be an effective measure for that purpose. However, such decision-making must be made with care because excessive replacement can trigger local extinctions. The barrier harvesting strategy moves any excess above a barrier to a different location. An appropriate selection of the barrier is thus important and the performance index to be obtained below will provide essential information for such decision making.

A (general) harvesting strategy can be modeled by a non-decreasing and right-continuous process *η* = (*η*_*t*_, *t* ≥ 0) with *η*_0_ = 0 (e.g., [47]). The random variable *η*_*t*_ for each *t* ≥ 0 denotes the aggregate harvested population artificially replaced from the storage until *t*. The process *η* needs to be adapted in the sense that the decision at time *t* must be made based only on the information available until *t*. Given a harvesting strategy *η*, the resulting storage population solves the SDE

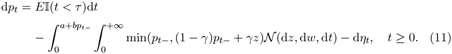

The barrier-type strategy, which we first focus on, is a practical harvesting strategy that continuously moves the excess above *P* of the storage population. Because the process *p* increases at a constant rate *E*, the strategy pushes the process downward at rate *E* whenever *p* is at *P* (so that it stays at *P* until the next outflow). Therefore, the barrier-type strategy, say 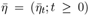, modeling the cumulative harvested population, is concisely given by 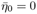 and

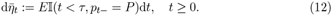

The resulting storage population is (11) with *η* replaced with 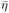.

We are particularly interested in the expected harvested fish population by the barrier-type strategy (12) until the completion time

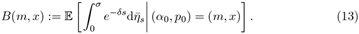

Here, we consider an exponentially discounted value, motivated by the following remark.

##### Remark 1

The discount factor exp(−*δs*) with a discount rate *δ* ≥ 0 is introduced here to account for the mortality of individual fish, namely shrinkage of the population, in the inflow, storage, and outflow processes at the same rate, so that it does not violate the mass conservation law (1). This formulation can effectively consider the exponential population decay (e.g., Chapter 1 of [55]), and moreover guarantees the uniqueness of numerical solutions to the associated BKE.

#### 3.1.3. Harvesting by a threshold-type strategy

Another important harvesting strategy considered in this paper is the threshold-type strategy that harvests *the outflow population rather than the storage population*. This strategy has the same motivation with the barrier-type strategy, aiming at sustaining the fish population in an anthropogenically disturbed river system by catching and transporting the outflow fish population. In practice, the threshold-type strategy can be achieved by casting a net at a location in the upstream side of a weir to capture the migrating fish in the outflow population (see, e.g., [52]).

Again, because this strategy controls the outflow instead of the storage population, it is mathematically distinct from the barrier-type strategy defined in Section 3.1.2. Importantly, it harvests only at discrete times 𝒯 (see (6)) when the outflow process *q* increases. This is in contrast to the barrier-strategy that modifies continuously the storage population *p*.

We formally define the threshold-type strategy as follows. At each jump time *t* in 𝒯, it harvests a fixed ratio *u* ∈ [0, 1] of the outflow. We also allow it to be capped at some *U* ∈ (0, ∞] to incorporate realistic technical capacities (e.g., net size). Thus, the harvested amount becomes min{*U, u*∆*q*_*t*_}. The uncapped version *U* = ∞ can also be considered, as a special case. The expected harvested amount of the fish population until completion is, for all (*m, x*) ∈ ℳ_0_ × [0, ∞),

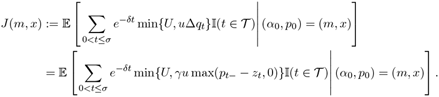

The discount factor exp(−*δt*) with *δ* ≥ 0 is again introduced to effectively account for the mortality (see Remark 1).

### 3.2. Governing BKEs

We now present the BKE governing the performance indices introduced in the previous subsection, written in terms of the associated infinitesimal generator.

The state process (*α, p*) is a bivariate Markov process on ℳ_0_×[0, ∞), with dynamics given by (3) and (7). The state space is decomposed as ℳ_0_ × [0, ∞) = *D*_0_ ∪ {(0, 0)} where

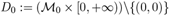

is transient and (0, 0) is absorbing.

The infinitesimal generator for (*α, p*) conditioned on (*α*_*t*_, *p*_*t*_) = (*m, x*) ∈ ℳ_0_ ×[0, ∞) for a generic sufficiently regular function Φ : ℳ_0_ ×[0, ∞) → ℝ is given by, for (*m, x*) ∈ *D*_0_,

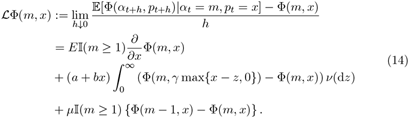

In particular, for the case *x* = 0,

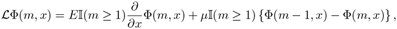

where the partial derivative in the right-hand side is understood to be the right-hand derivative. The boundary *x* = 0 is reflexive in the sense that *p* leaves 0 instantaneously because the population is continuously pushed toward the interior of *D*_0_ as long as *m* ∈ ℳ. This observation applies to the other BKEs presented in what follows.

#### 3.2.1. BKE for T, V and J

The governing BKE of the expectation of the completion time *T* = *T* (*m, x*) as in (10) is given by

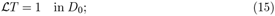

see, e.g., [56]. This also satisfies the boundary condition *T* (0, 0) = 0, which holds directly from the definition of *σ* in (9).

Similarly, we can obtain higher-order moments of the completion time by solving a cascading system of BKEs. In particular, the second-order moment *V* = *V* (*m, x*) is found from

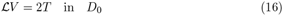

using *T* which can be obtained by (15). The boundary condition is *V* (0, 0) = 0.

The governing BKE of the performance index *J* (*m, x*) of the threshold-type strategy is given by

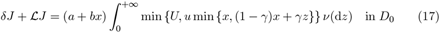

subject to the boundary condition *J* (0, 0) = 0.

#### 3.2.2. BKE for B

In a similar fashion, the governing BKE of *B* = *B*(*m, x*) can be written in terms of ℒ. A major difference is that under the barrier-type strategy, the storage population *p* only takes values on [0, *P*]. The controlled process (*α, p*) defined by (11) is a Markov process on ℳ_0_ × [0, *P*], which can be decomposed as {(0, 0)} ∪ *D*_*P*,1_ ∪ *D*_*P*,2_ where

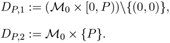

Here, *D*_*P*,1_ and *D*_*P*,2_ can be seen as waiting and controlling regions. Again (0, 0) is absorbing.

We have

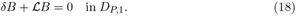

It also satisfies the (Dirichlet) boundary condition *B*(0, 0) = 0 as well as the Neumann boundary condition

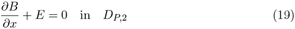

where the partial derivative in (19) is understood to be the left-hand derivative. Collectively, the function *B* can be seen as the solution of a mixed Dirichlet-Neumann boundary value problem (18)-(19).

### 3.3. Finite difference method

The BKEs (15), (16), (17), and (18) are written in terms of the common infinitesimal generator L. Hence, these equations can be numerically computed by using a common discretization scheme with minor modifications adapted to boundary conditions and additional terms. We propose a finite difference method to numerically discretize the BKEs and approximate the solutions. We firstly illustrate the discretization scheme for the BKEs for *T*, *V* and *J* and then that for *B*.

The discretized terms are assembled to find the numerical solution to each BKE. This is carried out by using a basic fast-sweeping method to numerically handle the assembled discretized equation (e.g., Section 4 of [57]).

#### 3.3.1. Finite difference method for T, V and J

For (15), (16) and (17), for which the state space ℳ_0_ × [0, ∞) is unbounded, we first truncate it to ℳ_0_ × [0, *K*] for large *K*. We divide [0, *K*] into *I* equally-spaced intervals and set *x*_0_ := 0, *x*_1_ := ∆*x, x*_2_ := 2∆*x*, …, *x*_*I*_ := *I*∆*x* = *K* where ∆*x* := *K/I*. Let ℐ := {0, 1, …, *I*} and define the set of grid points

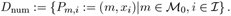

We aim to obtain approximation Φ_*m,i*_ ∈ *D*_num_ at the grid point *P*_*m,i*_ so that

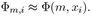

Each term of the infinitesimal generator (14) is discretized as follows.

1. The partial derivative 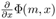 is approximated using the forward difference:

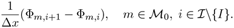
2. We approximate 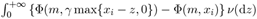 with

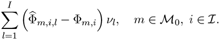 Here, 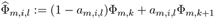 is a linearly interpolated value of Φ_*m,k*_ and Φ_*m,k*+1_ for the unique index *k* such that *γ* max {*x*_*i−l*_, 0} ∈ [*x*_*k*_, *x*_*k*+1_) and *a*_*m,i,l*_ := (*γ* max {*x*_*i−l*_, 0} − *x*_*k*_)*/*∆*x*. In addition, *ν*_*l*_ is a suitable approximation of 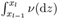.
3. Similarly, 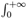 min {*U, u* min {*x*, (1 − *γ*)*x* + *γz*}} *ν*(d*z*) is approximated by

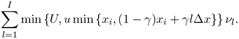

We thus obtain an approximation of (14): ℒΦ(*m, x*_*i*_) ≈ 𝒢 (*m, i*) where, for *i < I*,

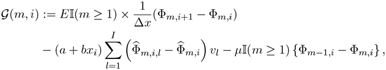

and for *i* = *I*,

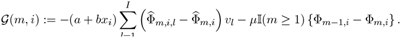

- For the cases Φ = *T* or Φ = *V*, the computation of Φ_*m,i*_ can be carried out by replacing ℑ*T* and ℑ*V* with 𝒢 in (15) and (16), respectively, and the boundary conditions Φ_0,0_ = 0.
- For the case Φ = *J*, the BKE (17) can be approximated by solving

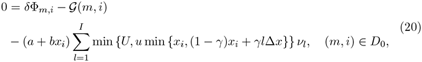

and the boundary condition Φ_0,0_ = 0.

#### 3.3.2. Finite difference method for B

For the computation of *B*, which is a solution to (18)-(19), because the state space ℳ_0_ × [0, *P*] is bounded, we can set *K* = *P* (without any truncation error). The BKE (18) can be approximated by

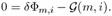

The Neumann boundary condition (19) is applied at the points *P*_*m,I*_, *m* ∈ ℳ and approximated as

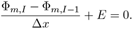

### 3.4. Monte-Carlo method

We conclude this section by presenting a time-explicit Monte-Carlo method to numerically generate sample paths of the population dynamics. This will be used to validate the finite difference method for the BKE in Section 4.

We fix a set of discrete times *t*_*n*_ := *n*∆*t* (*n* = 0, 1, 2, …) for small time-step ∆*t >* 0. Below, we illustrate a way to sample a path (*α*^(*n*)^, *p*^(*n*)^; *n* = 0, 1, 2, …) of the Markov process 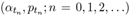. We focus on the case when there is no harvesting; the case with harvesting can be handled by straightforward modification.

First, we set *α*^(0)^ = *M*. When *α*_*t*_ = *m*, the transition time to *m* − 1 is exponential with parameter *µ*, which can be approximated by a geometric distribution. Hence, for each *n* ≥ 0, given *α*^(*n*)^ = *m*, we sample *α*^(*n*+1)^ so that it takes *m* − 1 with probability 1 − exp(−*µ*∆*t*) and *m* otherwise.

Regarding *p*, we have

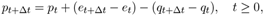

where

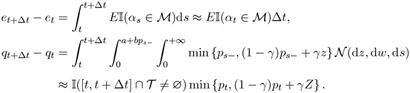

In the latter, the indicator is for the event the jump occurs on [*t, t* +∆*t*], which can be approximated by a Bernoulli random variable with parameter 1 −exp(−(*a* +*bp*^(*n*)^)∆*t*). The random variable *Z* is the first jump size after *t*, which is distributed according to *ν*.

Consequently, we set *p*^(0)^ = 0 and repeat

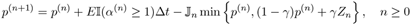

where we sample independent 𝕁_*n*_ ∼ Bernoulli(1 − exp(−(*a* + *bp*^(*n*)^)∆*t*)) and *Z*_*n*_ ∼ *ν*.

With this recursion, the non-negativity of each *p*^(*n*)^ is guaranteed due to

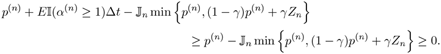

## 4. Applications

### 4.1. Target species

The target migratory fish specie we analyze in this section is the juvenile fish *P. altivelis*. This fish is diadromous and possesses a unique life span of one year. They migrate twice in their life times: spring upstream migration from the sea or a lake to a mid-stream river for growing and the autumn downstream migration from the midstream to the downstream of the river for spawning (see e.g., [58, 59]). In our study, we focus on the spring-season upstream migration due to the data availability. The study site is the Meiji-yousui weir, which has been utilized for agricultural, domestic, and industrial use [60] located at the midstream Yahagi River, Aichi Prefecture, Japan. The stream of Yahagi River with the length of 118 (km) and the catchment area of 1, 830 (km^2^) starts from the Mount Owari with the height of 1, 908 (m) and flows into the Mikawa Bay [61]. The daily migrating populations from the downstream side to the upstream side of the weir in the spring migration (∆*q*_*t*_ in our model) have been observed in recent years and had been made available by [49, 62].

### 4.2. Parameter estimation

We calibrate the model parameters of the migration of *P. altivelis* at Meiji-yousui weir between 2010 and 2022. The data from 2010 to 2020 were taken from [49] and those in 2021 and 2022 from the open database of [62].

Table 1 summarizes the start and end dates of migration as well as the duration each year. Here, the start (resp., end) day corresponds to the first (resp., last) day on which the population count was positive during the observation period. In each year, the migration event started in April and ended in June or July. The empirical mean and standard deviation of the duration from the start to the end day over these years are 83.8 (day) and 16.5 (day), respectively.

**Table 1.** The start, end, and duration of the migration event in each year from 2010-2022 (Source: [49, 62]). Duration is computed as Start-End+1.

| Year | Start | End | Duration | Year | Start | End | Duration |
| --- | --- | --- | --- | --- | --- | --- | --- |
| 2010 | Apr 16 | Jul 15 | 91 | 2017 | Apr 12 | Jun 30 | 80 |
| 2011 | Apr 10 | Jul 15 | 97 | 2018 | Apr 12 | Jun 30 | 80 |
| 2012 | Apr 13 | Jul 12 | 91 | 2019 | Apr 17 | Jun 9 | 54 |
| 2013 | Mar 30 | Jul 15 | 108 | 2020 | Apr 7 | Jun 29 | 84 |
| 2014 | Apr 15 | Jul 11 | 88 | 2021 | Apr 6 | May 15 | 40 |
| 2015 | Apr 12 | Jul 9 | 89 | 2022 | Apr 2 | Jun 30 | 90 |
| 2016 | Apr 11 | Jul 10 | 91 |  |  |  |  |

The total count of the migrated population, *q*_*σ*_ in our model, varies tremendously each year, as summarized in Table 2. It ranges from 447,134 in 2019 to 10,030,840 in 2016 (the ratio between these figures amounts to 22.4). Table 3 shows the average, standard deviation, skewness, and kurtosis of the daily counts of the outflow population in each year, calculated based on the data made available in [49, 62]. For all these years, the data show high variance (standard deviation is higher than the average). Furthermore, the daily counts of the outflow population are characterized by the positive skewness and kurtosis. Such statistics can be suitably modeled by positive and intermittent jumps as we will verify numerically.

**Table 2.** The total counts of the outflow population from 2010-2022 (Source: [49, 62]).

| Year | Total Count | Year | Total Count |
| --- | --- | --- | --- |
| 2010 | 487,951 | 2017 | 1,440,609 |
| 2011 | 985,637 | 2018 | 2,307,520 |
| 2012 | 761,990 | 2019 | 447,134 |
| 2013 | 839,587 | 2020 | 1,103,486 |
| 2014 | 601,147 | 2021 | 603,673 |
| 2015 | 1,276,048 | 2022 | 913,896 |
| 2016 | 10,030,840 |  |  |

**Table 3.** Average, standard deviation, skewness, and kurtosis of the daily counts of the outflow population in each year (Source: [49, 62]).

| Year | Average | Standard deviation | Skewness | Kurtosis |
| --- | --- | --- | --- | --- |
| 2010 | 5.483.E+03 | 1.117.E+04 | 3.350.E+00 | 1.215.E+01 |
| 2011 | 1.038.E+04 | 1.887.E+04 | 3.287.E+00 | 1.255.E+01 |
| 2012 | 8.374.E+03 | 1.376.E+04 | 2.304.E+00 | 4.938.E+00 |
| 2013 | 7.774.E+03 | 1.384.E+04 | 4.618.E+00 | 3.041.E+01 |
| 2014 | 6.910.E+03 | 1.306.E+04 | 3.448.E+00 | 1.559.E+01 |
| 2015 | 1.484.E+04 | 2.761.E+04 | 2.855.E+00 | 8.741.E+00 |
| 2016 | 1.102.E+05 | 1.651.E+05 | 1.567.E+00 | 1.694.E+00 |
| 2017 | 1.801.E+04 | 2.742.E+04 | 2.550.E+00 | 7.373.E+00 |
| 2018 | 3.077.E+04 | 5.760.E+04 | 2.981.E+00 | 1.093.E+01 |
| 2019 | 8.436.E+03 | 1.044.E+04 | 2.339.E+00 | 6.572.E+00 |
| 2020 | 1.314.E+04 | 1.873.E+04 | 1.854.E+00 | 3.440.E+00 |
| 2021 | 1.548.E+04 | 3.046.E+04 | 2.927.E+00 | 1.073.E+01 |
| 2022 | 1.039.E+04 | 1.940.E+04 | 2.901.E+00 | 9.115.E+00 |

We observe that the total migration count and the duration are only weakly correlated. The correlation coefficient between the duration in Table 1 and the total counts in Table 2 is 0.157. In other words, the influences of the total inflow on the completion time is small. In view of this large fluctuation of the population size with small correlation with the duration, we normalize the population dynamics with respect to the inflow rate *E* noting that the total counts are approximately proportional to *E* if the mortality is ignored. The normalization of (5) with respect to *E* leads to the rescaled dynamics:

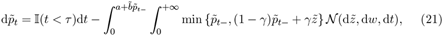

for *t* ≥ 0, with the scaled stored population 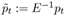 and scaled parameters 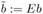 and 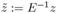. Here the time *t* is not scaled. Hereafter, we discuss the scaled dynamics (21) and omit tilde from these notations for simplicity. We assume an exponential-type jumps with *ν*(*x*) = *λe*^*−λx*^, *x* ≥ 0, for the Poisson random measure 𝒩.

The model parameters were estimated through a trial-and-error approach so that the obtained average (Ave) and standard deviation (Std) of the completion times have a minimal squared error with the empirical values: Ave*T*_*EM*_ := 83.8 (day) and Std*T*_*EM*_ := 16.5 (day). More precisely, we choose a parameter set with corresponding average Ave*T* and standard deviation Std*T* so that Err := (Ave*T* − Ave*T*_*EM*_)^2^ + (Std*T* − Std*T*_*EM*_)^2^ is minimized. Here, for the computation of Ave*T* and Std*T*, we use Monte-Carlo simulation based on 300,000 sample paths with the time step 0.025 (day) and the total length of the computational period 150 (day) (6,000 time steps). We are interested in the self-exciting behaviors where *a* is small (relative to *b*), and found that the following parameter set serves well in minimizing Err as 0.0244 and Ave*T* = 83.9 (day) and Std*T* = 16.8 (day): *M* = 25, *µ* = 0.30 (1/day), *a* = 1 (1/day), *b* = 9 (1/day), *λ* = 1.0 (1/day), and *γ* = 0.5. Figure 3 shows a sample path of the identified model.

**Figure 3.**
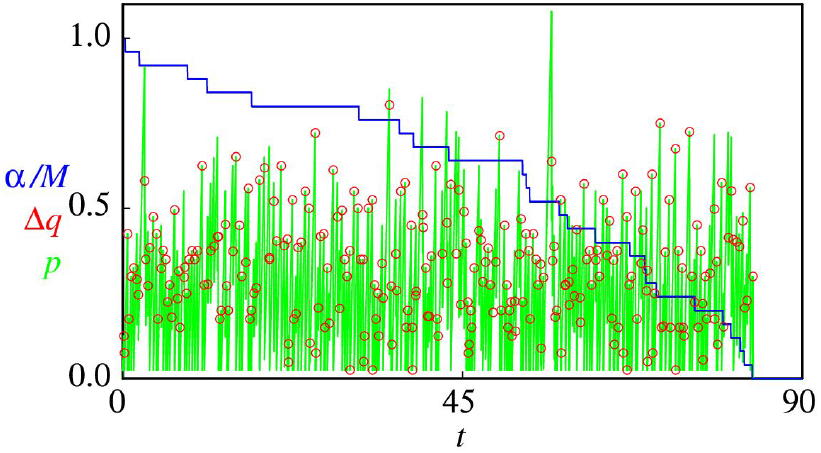
A sample path of the population dynamics with the identified model: *α*_*t*_*/M* (blue), ∆*q*_*t*_ (red), and *p*_*t*_ (green). The completion time of this path was computed as 83.475 (day). Here, ∆*q*_*t*_ is plotted only when it is positive.

We also examine the consistency between the Monte-Carlo method based on sample paths and the finite difference method based on the BKEs (15) for *T* and (16) for *V*. For the latter, we truncated with *K* = 5. Table 4 summarizes the average and standard deviation of the completion time of migration computed by these two methods, along with the empirical values. Overall, the produced values agree well, even with the largest considered ∆*x*(= *K/*200). Thus, for the rest of this section, we only provide the results computed by the finite-difference method.

**Table 4.** The mean (day) and standard deviation (day) of the completion time of migration computed by Monte-Carlo and BKE (finite difference) methods, along with empirical results.

|  |  | Average | Standard deviation |
| --- | --- | --- | --- |
| Empirical |  | 83.8 | 16.5 |
| Monte-Carlo (Sample paths) |  | 83.9 | 16.8 |
| Finite difference with $K = 5$ (BKE) | $\Delta x = K/1600$ | 83.9 | 16.7 |
| | $\Delta x = K/800$ | 83.9 | 16.7 |
| | $\Delta x = K/400$ | 83.8 | 16.7 |
| | $\Delta x = K/200$ | 83.8 | 16.7 |

### 4.3. Mean completion time

We analyze the completion time *σ* using the finite difference method based on the BKEs given in (15) and (16). We fix the step size of the finite difference method as ∆*x* = *K/*400 with *K* = 5, which gives a sufficiently fine grid in view of the results in Table 4.

Figure 4 shows the computed average completion time *T* = *T* (*m, x*) for the identified parameter values (see Section 4.2). Thanks to the use of the monotone finite difference method, the profile of the numerical solution is free from spurious oscillation. For *m* = 0, the obtained curve *x* ↦ *T* (0, *x*) starts at the origin and instantaneously jumps upward. It then decreases sharply for small *x* and tends to increase slowly for large *x*. This is an important behavior realized by the self-exciting feature we incorporated in the model (in the absence of self-exciting factor, the curve would be monotonically increasing). This unimodal behavior can be observed more clearly with different parameters (see also Figure 11). On the other hand, when *m* = 25, it is almost invariant to the value of *x*. In Figure 5, we show the computed standard deviation 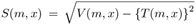. Similar to what we observed for the mean in Figure 4, for *m* = 0, it jumps at *x* = 0 and decreases on (0, ∞), whereas, when *m* = 25, the dependence on *x* is negligibly small.

**Figure 4.**
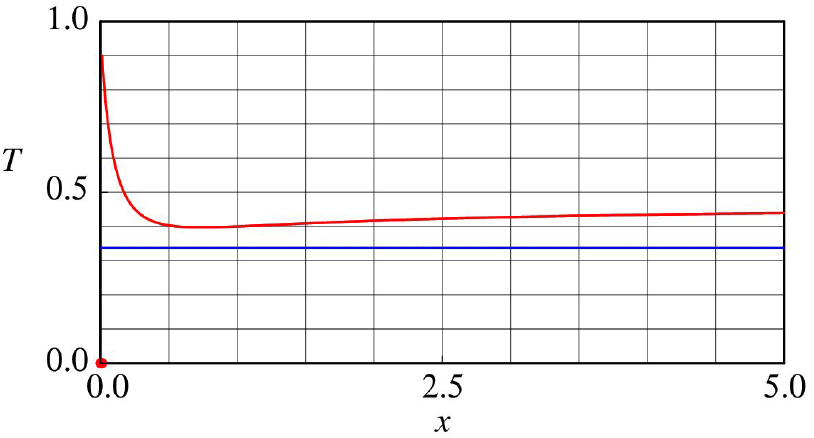
Computed mean completion time *T* = *T* (*m, x*) (day) for the identified parameter value: *m* = 0 (red) and *m* = 25 (blue). The value of *T* (25, *x*) is shifted by 83.5 downward simply for the illustration purpose. The red curve is discontinuous at the origin *x* = 0 and *T* (0, 0) = 0.

**Figure 5.**
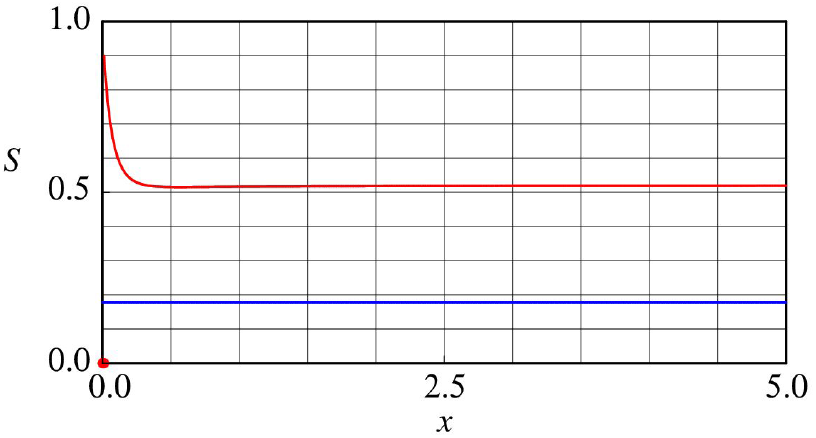
Computed standard deviation of the completion time *S* = *S*(*m, x*) (day) for the identified parameter value: *m* = 0 (red) and *m* = 25 (blue). The value of *S*(25, *x*) is shifted by 16.5 downward simply for the illustration purpose. The red curve is discontinuous at the origin *x* = 0 and *S*(0, 0) = 0.

We now perform sensitivity analysis of the mean completion time *T* with respect to the parameters *a, γ* and *λ*. The parameters *a* and *γ* regulate the degree of self-excitement in the population dynamics (see (4)). The parameter *λ* controls the jump size.

Figures 6–8 show the computed mean completion time *T* for different values of *a, γ* and *λ*, respectively, all as functions of *x*. We first consider the case *m* = 0 to see the impact of the parameter clearly (in view of the results obtained in Figures 4 and 5). The results for *m* = 25 are given subsequently.

**Figure 6.**
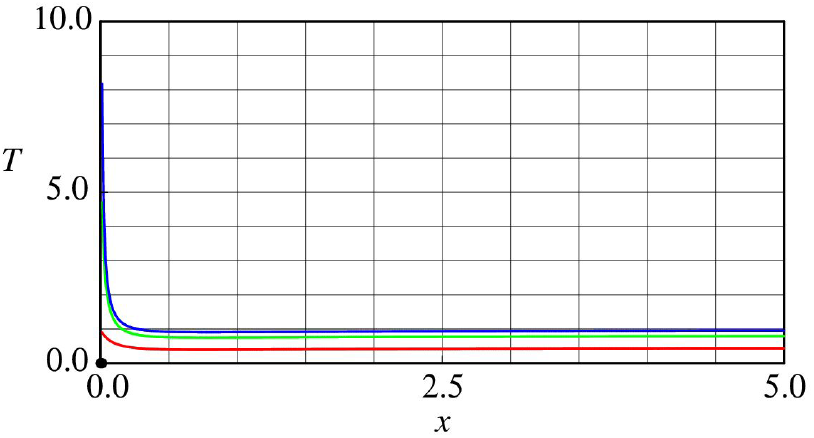
Computed mean completion time *T* = *T* (0, *x*) (day): *a* = 1 (1/day) (red), *a* = 0.1 (1/day) (green), *a* = 0.01 (1/day) (blue). All curves are discontinuous at the origin *x* = 0 and *T* (0, 0) = 0.

As shown in Figure 6, decreasing *a* leads to not only a larger mean completion time (via the resulting decrease in the net jump intensity of the outflow), but also a sharper spike around the origin *x* = 0. This computational result suggests that a sufficient amount of time, nearly 10 (day), is necessary for the population to complete migration when the storage population becomes near zero (by negative jumps).

Figure 7 shows that the change in parameter *γ* does not affect the profile of *T* (0, *x*) near the origin *x* = 0, but it does for *x* away from 0 in a non-monotone manner. Overall, *T* (0, *x*) is monotonically increasing in *γ*. This suggests that keeping the exogenic disturbance to stipulate the migration, such as the hydrodynamic disturbance, is crucial for sustaining the migration toward the upstream of the weir. In particular, during the migration period of the fish, one needs to prevent small flow velocity, caused by water infrastructures such as dams and weirs. A longer sojourn time in the storage may increase the risk of attacks by predators, such as piscivorous waterfowl.

**Figure 7.**
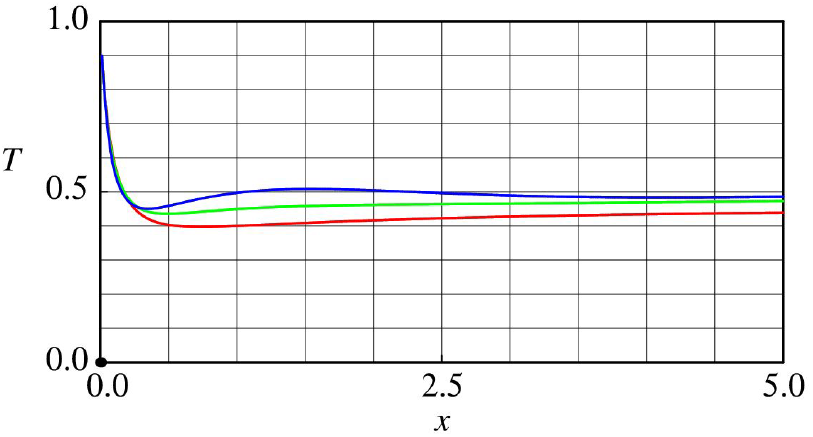
Computed mean completion time *T* = *T* (0, *x*) (day): *γ* = 0.5 (red), *γ* = 0.25 (green), *γ* = 0.125 (blue). All curves are discontinuous at the origin *x* = 0 and *T* (0, 0) = 0.

Figure 8 shows the computed *T* (0, *x*) for different values of *λ* (parameter for the jump size). For each *x >* 0, it is monotonically increasing in *λ* because jumps tend to be small for large *λ*. For each *λ*, as in the case of Figure 4, the curve *x* ↦ *T* (0, *x*) initially decreases and then increases. It is noted that it increases more sharply when *λ* is large. In addition, the minimizer of *x* ↦ *T* (0, *x*) tends to decrease in *λ*.

**Figure 8.**
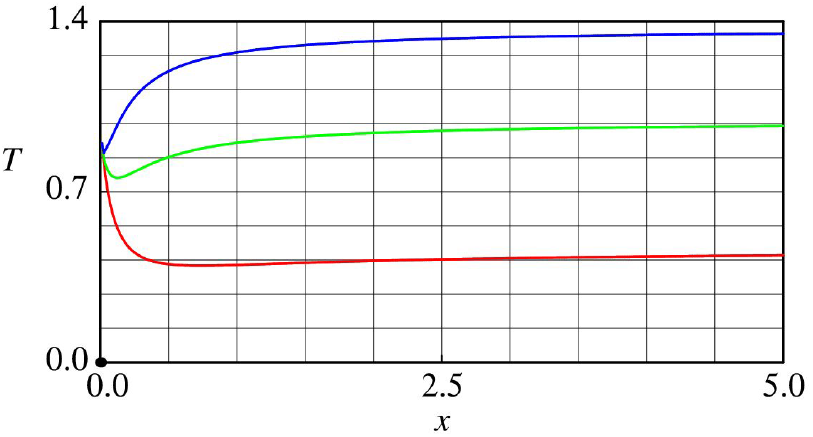
Computed mean completion time *T* = *T* (0, *x*) (day): *λ* = 1 (1/day) (red), *λ* = 4 (1/day) (green), *λ* = 8 (1/day) (blue). All curves are discontinuous at the origin *x* = 0 and *T* (0, 0) = 0.

We now analyze the case *m* = *M* = 25 to understand the impacts of (*a, γ, λ*) on the whole migration from the start to the end of the process. Figures 9 and 10 show the computed mean *T* (25, 0) and variance *S*(25, 0), respectively, as functions of *a* (1/day), *γ* and *λ*. In Figure 9, the mean *T* (25, 0) is decreasing with respect to *a* and is increasing with respect to *λ*. These are consistent with what we observed in Figures 6 and 8. By contrast, *T* (25, 0) is insensitive to the change in *γ* for the range examined, although this parameter affects the stored population dynamics after terminating the inflow as demonstrated in Figure 7. In view of Figure 10, we see that the standard deviation *S*(25, 0) is only sensitive to *a* (especially when it is small). It is almost invariant to the change in *γ* and *λ*.

**Figure 9.**
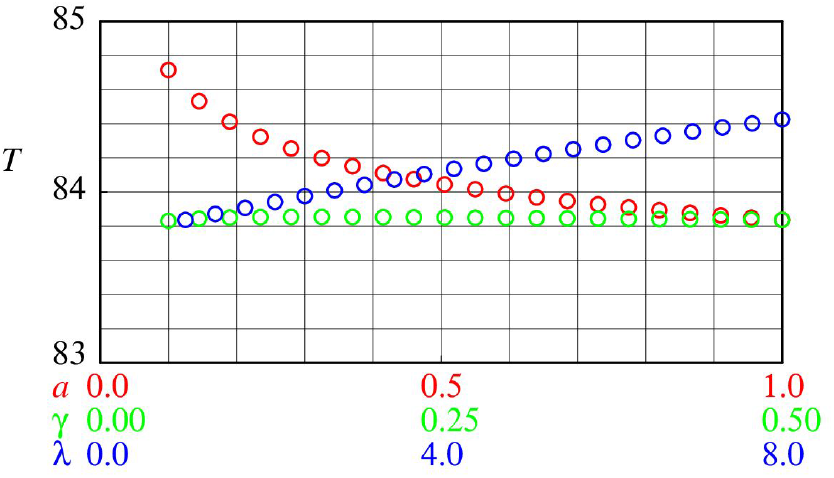
Computed mean completion time *T* = *T* (25, 0) (day) of the whole migration process for different values of *a* (1/day), *γ*, and *λ*.

**Figure 10.**
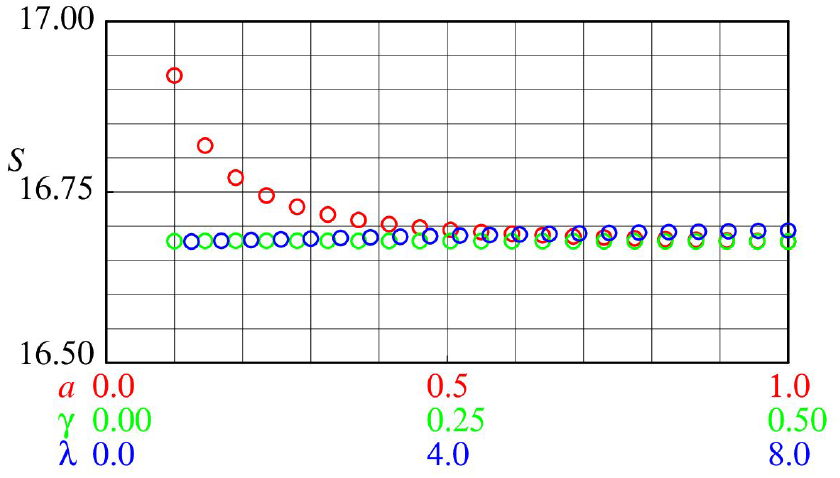
Computed standard deviation *S* = *S*(25, 0) (day) of the whole migration process for different values of *a* (1/day), *γ*, and *λ*.

**Figure 11.**
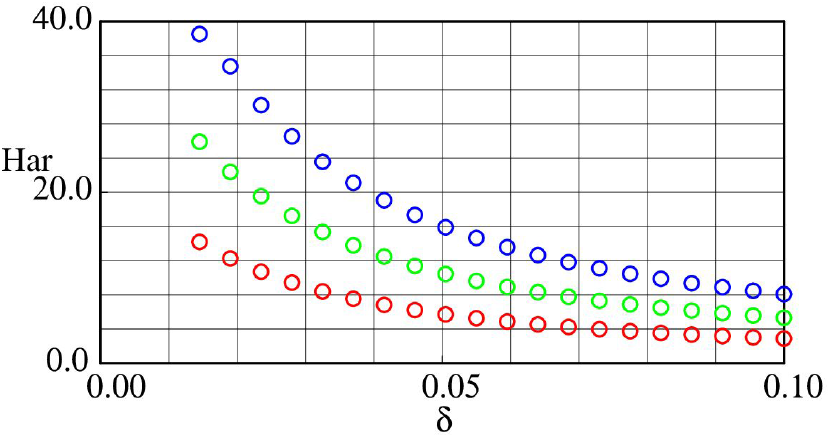
The harvested amount *B*(*M*, 0) under the barrier-type strategy as functions of *δ* (1/day) for *P* = 0.5 (red), *P* = 0.25 (green), and *P* = 0.125 (blue).

### 4.4. Partial harvesting problem

We compute the total harvested population under the barrier- and threshold-type strategies studied in Sections 3.1.2 and 3.1.3 and conduct sensitivity analysis. Here, the mean total inflow is *E*𝔼[*τ*] = 𝔼 [*τ*] = *M/µ* = 83.3 (recall that we assume the normalized case). Unless otherwise specified, we set the discount rate *δ* = 0.01 (1/day). For the barrier-type strategy we set the barrier height *P* = 0.5; for the threshold-type strategy we set the parameters *U* = 0.1 and *u* = 0.2. In each figure panel, “Har” corresponds to either the harvested amount *B*(*M*, 0) by the barrier-type strategy or *J* (*M*, 0) by the threshold-type strategy where *M* = 25. We use *K* = 5 for truncation for the threshold-type strategy.

Figures 11 and 12, respectively, show the harvested amount for different values of *P* and *u* under the barrier- and threshold-type strategies, as functions of *δ*. In both cases, the harvested amounts decrease sharply as the mortality *δ* increases. The harvested amount is decreasing in the barrier *P* under the barrier-type strategy, and also in the harvesting ratio *u* under the threshold-type strategy. These results are consistent with our intuition.

**Figure 12.**
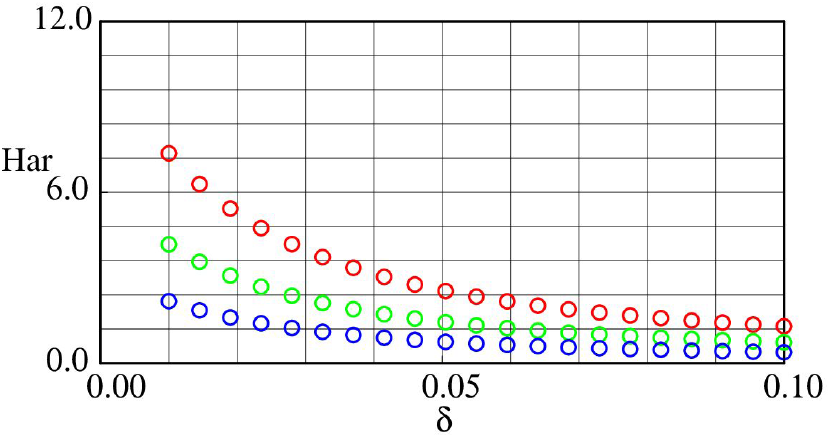
The harvested amount *J* (*M*, 0) under the threshold-type strategy as functions of *δ* (1/day) for *u* = 0.05 (red), *u* = 0.025 (green), and *u* = 0.0125 (blue).

Figures 13 and 14 show the harvested amounts under both strategies as functions of *a* and *γ*, respectively. In contrast to those shown in Figures 11-12, the dependence on *a* and *γ* are less intuitive, as we analyze below.

**Figure 13.**
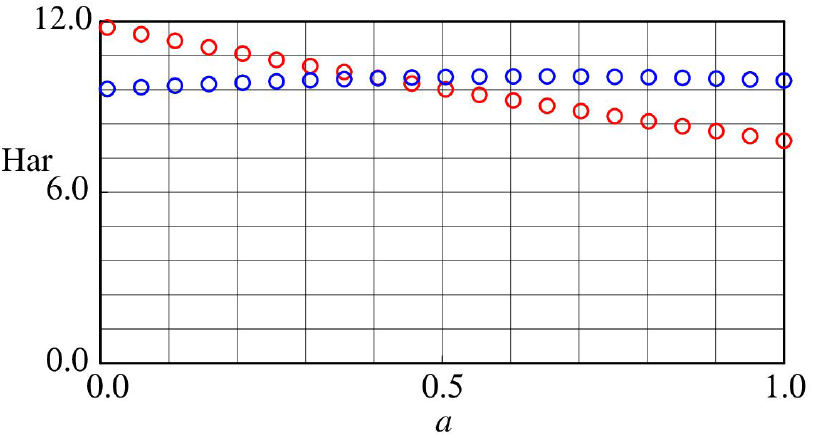
The harvested amount *B*(*M*, 0) under the barrier-type strategy (red) and *J* (*M*, 0) under the threshold-type strategy (blue) as functions of *a* (1/day).

**Figure 14.**
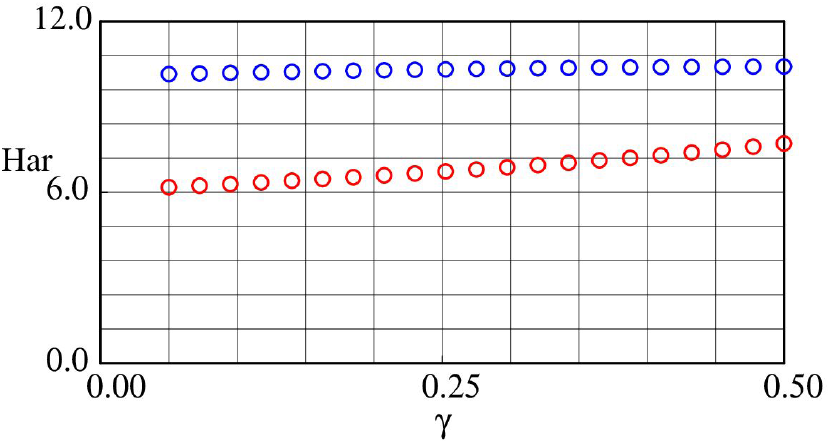
The harvested amount *B*(*M*, 0) under the barrier-type strategy (red) and *J* (*M*, 0) under the threshold-type strategy (blue) as functions of *γ*.

As can be observed in Figure 13, the harvested amount under the barrier-type strategy *B*(*M*, 0) decreases in the exogenic jump rate *a*. Note that harvesting occurs precisely when the stored population *p* is as high as *P*. The stored population at each time tends to decrease in *a* (see also Figure 6) and this explains why the harvested amount is decreasing. As for the threshold-type strategy, *J* (*M*, 0) depends non-monotonically on *a*, with its peak at around *a* ≈ 0.65. These results imply that there exists an optimal exogenic disturbance level (via modulating the flow discharge or its fluctuation, for example) that maximizes the harvested amount under the considered threshold-type strategy for *P. altivelis* at the study site. This non-monotone profile of *J* (*M*, 0) is mainly due to the capacity *U*. To see this, when *U* = ∞, the *u*-fraction of all fish that enter the storage will be eventually harvested. Hence, the harvested amount is invariant to *a* (if mortality is ignored). On the other hand, when *U* is finite, any excess above *U* (which depends on *a*) fails to be caught. Note that the value of *J* (*M*, 0) is not monotone in *a* because the discount/mortality at rate *δ* also plays a role.

In Figure 14, we observe that, under both the barrier- and threshold-type harvesting strategies, the harvested amount increases in *γ* (which represents the self-excitedness or memory in the outflow process). On the other hand, it is more sensitive in the change of *γ* for the barrier-type strategy in comparison to the threshold-type strategy.

In summary, our computational study discussed in this section demonstrated that the proposed mathematical model is consistent with the empirical data, and can be utilized for the evaluation of the harvesting strategies of *P. altivelis* and other fishes (see, e.g., [63, 64]) in the studied river environment.

## 5. Conclusion

This paper proposed a jump-driven self-exciting stochastic process model for describing the upstream fish migration and applied it to study the migration of *P. altivelis* with the existing record collected in a river in Japan. The model was formulated as a single-variable model and its associated BKEs were obtained along with the finite difference method for the numerical computation, which was confirmed to be consistent with the results produced by the Monte-Carlo method. The barrier- and threshold-type harvesting strategies were investigated numerically using the developed finite difference methods.

The proposed stochastic process model written in terms of SDEs and the developed computational schemes can be incorporated and/or adapted to study other fish migration events and also to solve optimization problems toward the conservation of migratory fish species. Examples of potential future research include the application in the barrier removal/restoration planning in river systems [65, 66]. Coupling the proposed model with a fish-schooling model [54, 67] to focus more on local migration behavior of fish populations is another venue for future research. Furthermore, our developed stochastic process model can be directly used to describe the population dynamics in the long-term resource management problem [68]. A more sophisticated model can be pursued by clarifying the interaction between fish migration and hydrological disturbance [69, 70], while the challenge will be the identification of their parametric relationship. Finally, the contribution of diadromous fish species to the global carbon transport [71] can be re-examined by using the proposed model.

